# RyR2 inhibition with dantrolene is antiarrhythmic, antifibrotic, and improves cardiac function in chronic ischemic heart disease

**DOI:** 10.1101/2022.06.24.496861

**Authors:** Jeffrey Schmeckpeper, Kyungsoo Kim, Sharon A George, Dan Blackwell, Jaclyn A Brennan, Igor R Efimov, Bjorn C Knollmann

**Affiliations:** Vanderbilt Center for Arrhythmia Research and Therapeutics, Division of Clinical Pharmacology, Vanderbilt University Medical Center, Nashville, Tennessee, USA; Department of Biomedical Engineering, the George Washington University, Washington DC; Department of Biomedical Engineering, Northwestern University, Chicago IL

**Keywords:** Dantrolene, Ryanodine Receptor, Ventricular Tachycardia, Heart Failure

## Abstract

**Background:** Ventricular tachycardia (VT) is responsible for sudden death in chronic ischemic heart disease (CIHD) patients. The cardiac ryanodine receptor (RyR2) releases Ca2+ from the sarcoplasmic reticulum (SR) and links electrical excitation to contraction. RyR2 hyperactivity has been widely documented in CIHD and may contribute to VT risk and progressive LV remodeling.

**Objective:** To test the hypothesis that targeting RyR2 hyperactivity plays a mechanistic role in VT inducibility and progressive heart failure in CIHD that can be prevented by the RyR2 inhibitor dantrolene.

**Methods:** CIHD was induced in C57BL/6J mice by left coronary artery ligation. Four weeks later, mice were randomized to either acute or chronic (6 weeks via osmotic mini-pump) treatment with dantrolene or vehicle. VT inducibility was assessed by programmed stimulation *in vivo* and in isolated hearts. Electrical substrate remodeling was assessed by optical mapping. Ca2+ sparks and spontaneous Ca2+ releases were measured in isolated cardiomyocytes. Cardiac remodeling was assessed by histology and qRT-PCR. Cardiac function and contractility were assessed by echocardiography.

**Results:** Compared to vehicle, acute dantrolene treatment reduced VT inducibility and improved LV contractility *in vivo*. Optical mapping in isolated hearts demonstrated reentrant VT prevention by dantrolene, which normalized the shortened refractory period (VERP) and prolonged action potential duration (APD), preventing APD alternans. In single CIHD cardiomyocytes, dantrolene normalized RyR2 hyperactivity and prevented spontaneous SR Ca^2+^ release. Chronic dantrolene treatment reduced peripheral muscle strength but had no adverse effects on body weight or mortality. Chronic dantrolene not only reduced VT inducibility but also reduced peri-infarct fibrosis and prevented the progression of LV dysfunction in CIHD mice.

**Conclusion:** RyR2 hyperactivity plays a mechanistic role for VT risk, infarct remodeling, and contractile dysfunction in CIHD mice. Our data provide proof of concept for the anti-arrhythmic and anti-fibrotic efficacy of dantrolene in CIHD.

**Clinical Perspective:** *What is New?:*

- The mouse CIHD model is a more clinically relevant model in which treatment is started late after infarction, when heart failure is already established.
- Acute and chronic dantrolene treatment suppresses VT inducibility by restoring myocyte APD, terminating APD alternans and normalizing VERP.
- Chronic dantrolene treatment prevents pathological remodeling and peri-infarct fibrosis, the substrate for reentry VT. Cardiac function is improved with chronic dantrolene therapy.

*Clinical Implications:*

- Treatment with dantrolene, which is already approved for clinical use, is a promising therapy in patients with ischemic heart disease, in whom other antiarrhythmic drugs are contraindicated.
- Dantrolene inhibition of RyR2 not only suppresses VT but also improves cardiac function in chronic ischemic heart disease.

## Introduction

Sudden cardiac death (SCD) due to ventricular tachycardia (VT) or ventricular fibrillation (VF) is a significant public health problem, accounting for up to 20% of all deaths in adults in the US.^1^ The vast majority (>90%) of SCD occurs in patients with coronary disease. While implantable cardioverter-defibrillators (ICDs) are the primary therapeutic option for cardiac arrest survivors after myocardial infarction and others at high risk for SCD, ICDs terminate arrhythmias after they occur and many patients with and without ICDs continue to require anti-arrhythmic drugs to prevent arrhythmias. Currently available anti-arrhythmic drugs targeting ion channels on the cell surface have limited efficacy when used acutely in patients with structural heart disease and can worsen heart failure. Their chronic use provides either no survival benefit or increases mortality due to pro-arrhythmic effects.^2, 3^ New therapeutic approaches are needed to prevent arrhythmia and SCD in patients with structural heart disease.

The cardiac ryanodine receptor 2 (RyR2) releases calcium (Ca2+) from the sarcoplasmic reticulum (SR) to coordinate cardiac excitation-contraction coupling. Dysfunction of RyR2 leads to SR Ca2+ leak during diastole, which reduces SR Ca2+ content and is energetically costly to a failing heart.^4^ Additionally, arrhythmogenic spontaneous Ca2+ release events in cardiomyocytes isolated from animal models of heart failure ^5^ have been related to an increase in the RyR2 phosphorylation status by PKA or Ca2+/calmodulin-dependent protein kinase II.^6, 7^ Mutations that render RyR2 hyperactive cause catecholaminergic polymorphic ventricular tachycardia (CPVT), where catecholamine-induced spontaneous Ca2+ release from SR via RyR2 generates potentially fatal cardiac arrhythmias. The current model proposes that these modifications lead to conformation changes in RyR2, which allows spontaneous diastolic SR Ca2+ release, resulting in delayed afterdepolarizations (DADs) and triggered beats that can generate ventricular tachyarrhythmias.^8^ Evidence from modeling and animal studies suggests that Ca2+ leak triggers ventricular ectopy and generates an arrhythmogenic substrate that can support monomorphic VT through multiple mechanisms. ^6, 9, 10^ Furthermore, a rise in intracellular Ca2+ due to RyR2 opening may activate small conductance calcium-activated potassium channels, shortening refractory periods and facilitating reentry and ventricular fibrillation.^11^ Hence, normalizing the RyR2 hyperactivity^12^ that results in Ca2+ leak can be considered a promising strategy for preventing ventricular arrhythmias associated with structural heart disease.

After myocardial infarction, the heart undergoes substantial remodeling associated with inflammation, replacement fibrosis, cardiomyocyte hypertrophy and chamber dilation driven by wall stress, cytokines and neurohormonal activation.^13^ Multiple animal models and human studies with ischemia and heart failure have shown that there is RyR2 hyperactivity due to oxidation and posttranslational modification, which causes Ca2+ leak in the setting of pathological remodeling.^14-19^ While the extent to which RyR2 modification contributes to left ventricular (LV) dysfunction in these models remains controversial^7^, it is clear that diastolic Ca2+ leak plays a role in decreased Ca2+ transients and reduced excitation-contraction coupling at the cellular level.^20^

Dantrolene has been used clinically for many years to suppress skeletal muscle Ca2+ leak due to mutations in RyR1 in malignant hyperthermia. Multiple studies have shown that dantrolene also stabilizes the tertiary structure of RyR2 and prevents diastolic Ca2+ leak in failing myocytes.^21^ Acute or short-term dantrolene treatment has been used to suppress RyR2 hyperactivity in induced ventricular and atrial arrhythmias^16, 22^, doxorubicin cardiotoxicity^23^, resuscitation models after ventricular fibrillation arrest^24^, and genetic models of RyR2 mutations that cause catecholamine polymorphic VT (CPVT).^25^ While long-term dantrolene treatment has shown promise in preventing ventricular arrhythmia, cardiac remodeling and reduced contractility in a model of tachycardia mediated heart failure^21^, to date, no study has evaluated long-term dantrolene treatment late after myocardial infarction and ischemic heart failure.

Here, we utilized an accepted murine model of chronic ischemic heart disease (CIHD) to provide proof of concept for the therapeutic efficacy of targeting RyR2 with dantrolene after myocardial infarction. Our experimental study demonstrates for the first time that preventing RyR2 hyperactivity not only suppresses DAD-triggered activity but also prevents reentrant VT induction in vivo. The anti-arrhythmic action of dantrolene is likely the result of improved SR Ca2+ handling, which normalized the shortened ventricular action potential and effective refractoriness that rendered CIHD hearts susceptible to reentrant VT. In addition, long-term treatment with dantrolene improved cardiac function and reduced fibrosis in the infarct border zone, which is a substrate for reentrant VT. Taken together, our results demonstrate that RyR2 hyperactivity not only contributes mechanistically to VT induction but also to adverse cardiac remodeling and progressive LV dysfunction in CIHD. RyR2 should be considered a therapeutic target for preventing VT and improving cardiac function in structural heart disease.

## Methods

### Mouse CIHD model

All studies were approved by the Vanderbilt Animal Care and Use Committee of Vanderbilt University, USA (Protocol M1900081-00) and performed in accordance with NIH guidelines. To induce CIHD, 10-12 week old male and female C57BL/6J mice underwent complete ligation of the left coronary artery as previously described.^26^ Mice were allowed to recover for 4 weeks before inclusion into the study. The inclusion criteria for CIHD mice were 1) Fractional shortening < 35%, 2) Ejection fraction <50%, and 3) mean velocity fiber shortening <2.5 circ/sec. A total of 39 male and 54 female mice were included in the study. All data analysis was performed in a blinded fashion regarding the treatment group.

For chronic dantrolene treatment, osmotic mini-pump (#2006, Alzet) were implanted subcutaneously in CIHD mice 4 weeks post coronary ligation per manufactures protocol. Dantrolene sodium suspension (Ryanodex, Eagle Pharmaceuticals) was diluted in 0.9% normal saline to deliver 20 mg/kg/day of dantrolene over 6 weeks.

### Mouse transesophageal programmed electrical stimulation (PES)

Sham and CIHD mice were anesthetized with inhaled isoflurane (3% for induction, 2-2.5% for maintenance) while breathing spontaneously and placed in the supine position on a heating pad. Surface ECG was recorded continuously using AD Instruments amplifiers and LabChart 8 software. An octopolar 2F electrode catheter (CIB’ER MOUSE™; NuMED, Inc) was placed in the esophagus via the mouth, guided by electrogram tracings to verify position. Unipolar pacing was performed using a programmable stimulator with 6 mA of pacing amplitude and 3 ms of pulse width for all studies. PES consisted of pacing with a train of 15 beats (10 Hz, S1), followed by a single extra stimulus (S2) to determine the ventricular effective refractory period (VERP). VT induction was then performed using 3 extra stimuli (S2-S4) following each pacing train to induce VT. Dantrolene (30 mg/kg, intraperitoneal injection) using Ryanodex (Eagle Pharmaceuticals, Inc., NJ) or 0.9% normal saline was administered to mice 30 minutes prior to the study. Isoproterenol (1.5 mg/kg, intraperitoneal injection) was administered after capturing ventricular pacing.

### Optical mapping of isolated mouse hearts

CIHD mice were anesthetized by isoflurane 4 weeks after coronary ligation. Hearts were isolated after thoracotomy and Langendorff perfused with a modified Tyrode’s solution (130 mM NaCl, 24 mM NaHCO3, 1.2 mM NaH2PO4, 4 mM KCl, 1 mM MgCl2, 5.6 mM Glucose and 1.8 mM CaCl2, pH 7.4 at 37°C). Cardiac motion was arrested during optical mapping using blebbistatin (15 μM). Hearts were paced using a bipolar platinum pacing wire placed on the anterior surface of the heart, at the center of the field of view, using pulses at 1.5x threshold of stimulation and 2 ms duration. Restitution properties were measured by pacing at multiple basic cycle lengths (BCL) from 200 - 60 ms. After a 15 min equilibration period, hearts were stained with di-4-ANEPPS, a voltage-sensitive dye (37.5 μg/ml in Tyrode solution). After a 5 min washout period, the dye was excited using light at 510±5 nm wavelength. Emitted fluorescence was filtered using a 610±20nm bandpass filter and recorded using a CMOS camera (MiCam05, SciMedia). Optical recordings were obtained at baseline, Isoproterenol (250 nM) treatment and Iso + Dantrolene (10 μM) treatment. Optical signals were analyzed using a custom Matlab program (Rhythm).^27^

### Action potential and Ca2+ transient measurements in isolated myocytes

Cardiomyocytes were loaded with Fura-2 acetoxymethyl ester (Fura-2 AM; Invitrogen) as described previously.^28^ After Fura-2 loading, experiments were conducted in NT solution containing 1 μM isoproterenol and 2 mM CaCl2. Fura-2 AM-loaded myocytes were pre-incubated for 1 hour with vehicle or 1 μM dantrolene. Spontaneous Ca2+ release events were quantified during the 30 seconds following cessation of the pacing train. Data were analyzed using IonWizard data analysis software (Milton, MA).

### Ca2+ sparks in permeabilized cardiomyocytes

Ca2+ sparks were measured in isolated adult myocytes from CIHD mice by spinning-disc confocal microscopy as previously described.^29^ Analysis of spark frequency was performed using the SparkMaster plugin for ImageJ and all data were normalized to SR Ca2+ content for the experimental day. Statistical comparisons were made using a linear mixed-effects (hierarchical) model^30^, clustered by mouse to account for random effects between isolations and calculate Bonferroni-adjusted p values.

### Statistics

Statistical analyses were performed using Prism v9.0.2 (GraphPad Software, Inc.) or R for linear regression modeling. Statistical tests used are reported in the statistical summary table (Supplemental Figure 1) and the figure legends. For normally distributed data (VERP, APD_80_, Infarct Size, Fibrosis area, FS, mcVcf, LVESD, LVEF), mean and standard deviation are provided. ANOVA with Bonferroni correction was used for multiple comparisons to assess for interaction between variables.For non-normally distributed data (VT episodes/train, VT duration, Ca2+ sparks, and Spontaneous Ca2+ release), the mean and 95% confidence intervals are provided. Linear regression models were used to assess the interaction between multiple variables and multiple comparisons when the data were non-normally distributed.

## Results

### CIHD renders mice susceptible to VT induction by programmed electrical stimulation

Programmed electrical stimulation (PES) is widely used to initiate sustained monomorphic VT and is traditionally considered a predictor of future arrhythmic events and mortality following myocardial infarction. We first tested whether PES could induce VT in CIHD mice. A transesophageal PES approach was used to allow for repeated measurement in the same mouse (Fig 1A). PES induced VT in 17.7 % of CIHD mice (6 of 34 mice: Fig 1C), whereas sham mice did not exhibit any inducible VT (Fig 1B&C). The addition of a catecholamine challenge with isoproterenol (PES+Iso) rendered 47.1% of CIHD mice inducible (16 of 34 mice: Fig 1C). There was no difference in VT induction between male and female mice (VT inducibility: male 4/9 vs female 12/25; P=.4709, Fischer Exact Test). Iso also significantly increased the frequency (Fig 1D) and duration of VT (Fig 1E) per CIHD mice. Taken together, these data demonstrate that PES+Iso reproducibly induces VT in CIHD mice.

**Figure 1.**
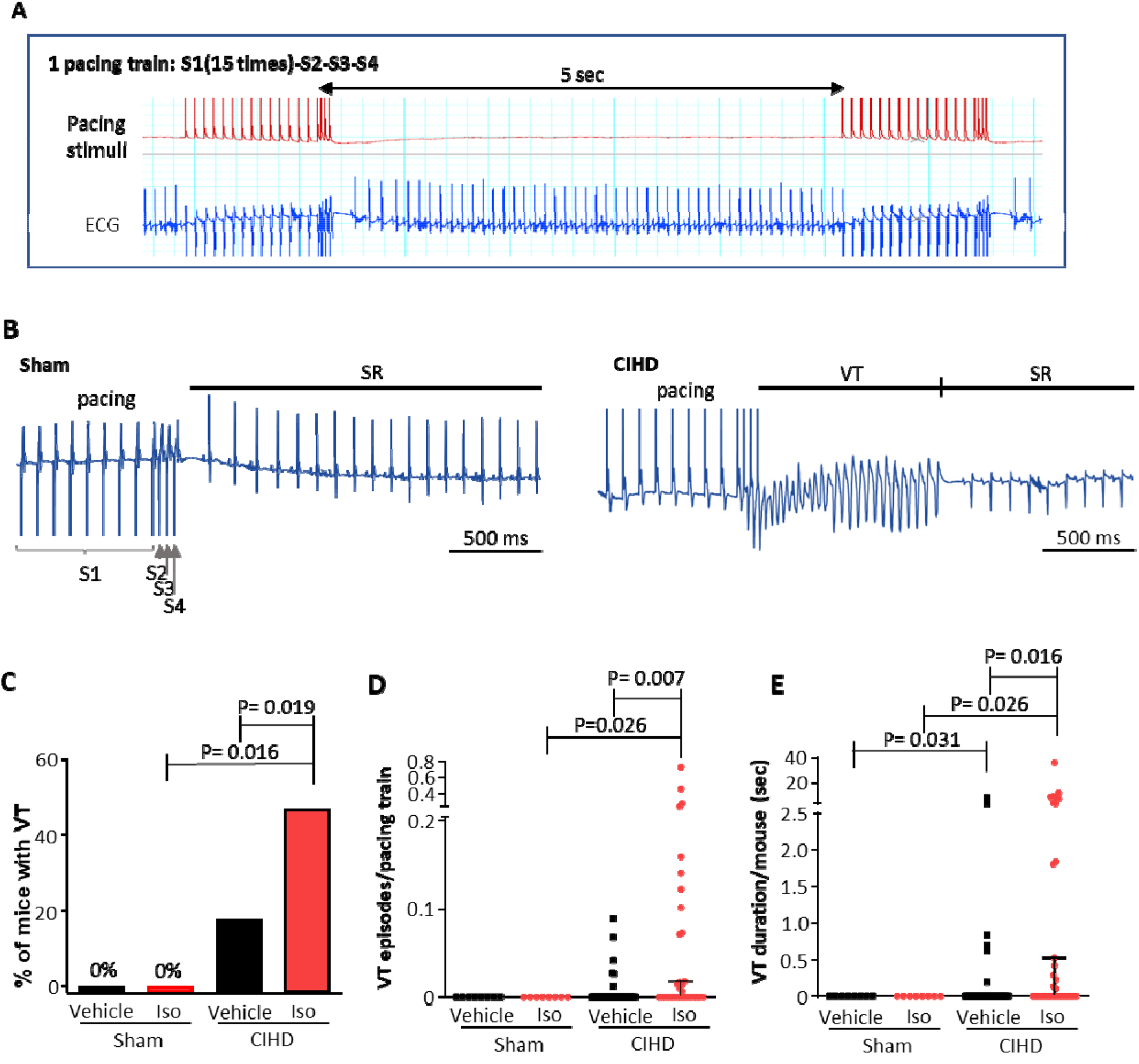
Inducibility of ventricular tachycardia (VT) by programmed electrical stimulation. (A) Stimulation protocol (top) and representative electrocardiogram (ECG, bottom). An episode of programmed stimulation protocol consists of 15 S1 stimuli followed by three (S2-S4) extra stimuli that are optimized with minimum intervals of ventricular capturing. (B) Representative ECG traces of sham (left) and CIHD (right) mice after an episode of pacing protocol in the presence of isoproterenol. CIHD mice exhibit inducible VT followed by spontaneous conversion to sinus rhythm (SR). (C) Incidence of inducible VT. P values were obtained using the Fischer Exact Test. (D) VT episodes/pacing train and (E) VT duration in sham and CIHD mice. P values were obtained using the Wilcoxon signed-rank test. [Sham N=8 and CIHD= 34 mice]

### Acute dantrolene treatment suppresses VT and PVC in CIHD

To test whether acute RyR2 inhibition with dantrolene can prevent VT induction, CIHD mice underwent a baseline PES followed by a PES in the presence of dantrolene one week later, followed by another PES 2 weeks later (Fig. 2A). Acute injection of dantrolene 30 min prior to the PES significantly reduced VT inducibility in CIHD mice by 33.3% (OR 0.0; P=0.039 Fischer Exact Test, Fig 2B). In the CIHD mice with VT, dantrolene also significantly reduced the frequency (Fig 2C) and duration of VT episodes (Fig 2D). Importantly, after the 2-week washout period, CIHD mice exhibited VT inducibility similar to baseline (Fig. 2B-E). Dantrolene also suppressed PVCs during PES+Iso, which returned to baseline numbers after washout (Fig2E). Together, these data demonstrate that dantrolene treatment suppresses both VT inducibility and PVCs in CIHD.

**Figure 2:**
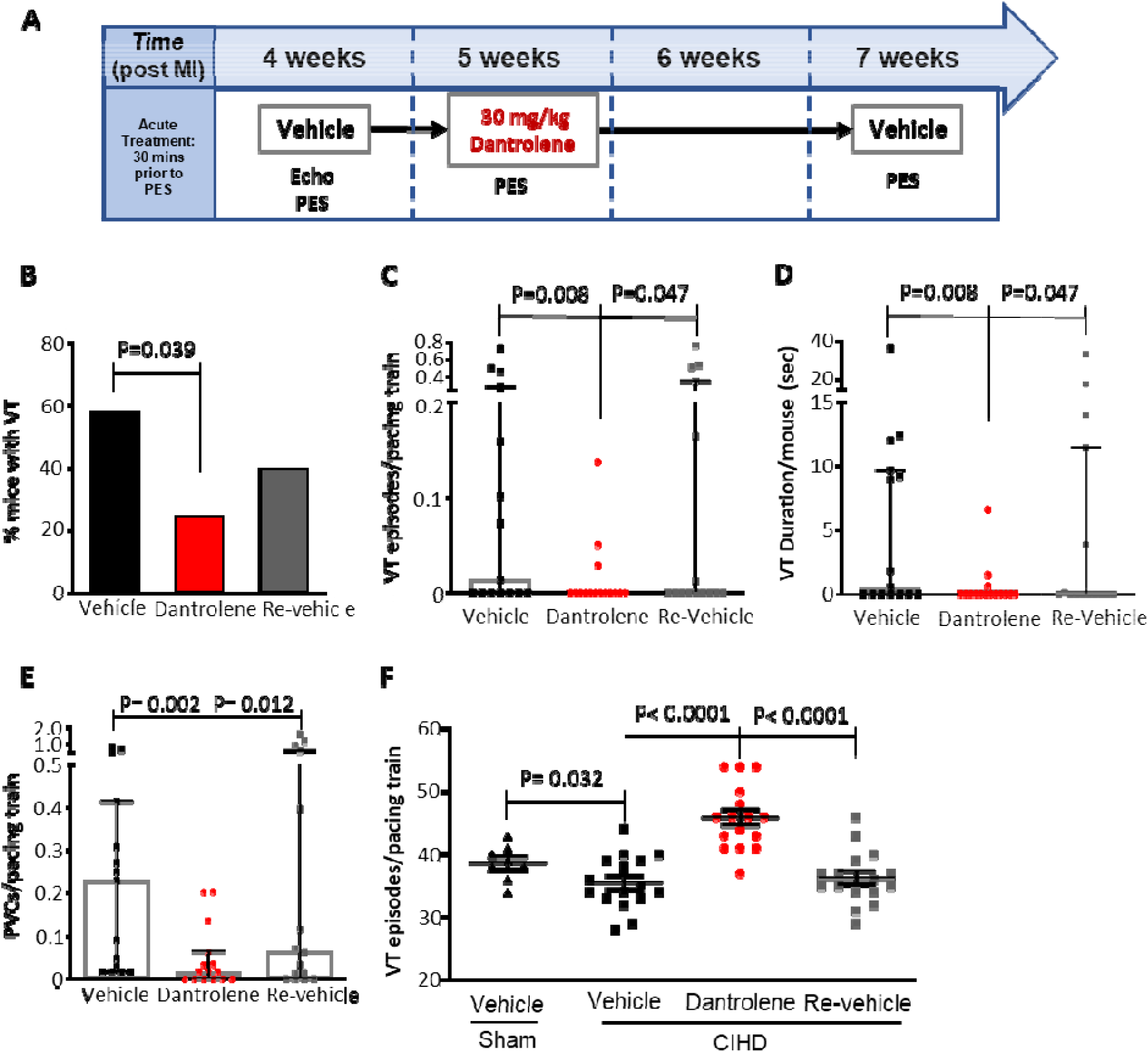
Acute dantrolene treatment reduces VT inducibility and ventricular ectopy in CIHD mice. (A) Experimental timeline. (B) Incidence of inducible VT by programmed electrical stimulation (PES). P values were obtained using the Fischer Exact Test. (C) VT episodes/pacing train and (D) VT duration per pacing train. (E) PVC frequency per pacing train. P values were obtained using the Wilcoxon signed-rank test. (F) Ventricular effective refractory period (VERP) measured during PES. P-values were obtained using Welsh ANOVA with Dunnett’s T3 multiple comparisons test. [N= 15 mice]

To determine why CIHD mice were susceptible to VT induction, we measured the ventricular effective refractory period (VERP). CIHD mice exhibit a shortened VERP (Sham 38.6±0.7 ms vs. CIHD 35.2±1.1 ms; P=0.032, Fig 2F), which can facilitate reentry arrhythmias. Acute dantrolene administration normalized the VERP of CIHD mice. After a 2-week washout, the VERP had returned to the short baseline values of CIHD mice. These data show that the normalization of the VERP is specific to dantrolene treatment rather than ongoing remodeling of the myocardium post-infarction.

### Acute dantrolene prevents reentry by suppressing APD alternans and prolonging APD in the infarct border zone

Optical mapping of action potentials was performed on isolated CIHD hearts to examine the mechanism of VT induction and its prevention by dantrolene. Burst pacing with Iso induced VT in all CIHD hearts and dantrolene treatment suppressed VT inducibility (Fig. 3D). Phase maps generated from optical action potentials demonstrated a reentrant pattern around the infarct scar (Fig. 3A) and VT induction (Fig. 3B) was observed in ECG and optical recordings. Consistent with its normalizing effect on VERP in vivo, dantrolene administration prolonged APD_80_ in isolated CIHD hearts (Fig 3C and E). Conduction velocity (CV) was not altered in CIHD hearts or affected by dantrolene treatment (Fig 3C and F). APD alternans, a known risk indicator for VT induction, were observed in isolated CIHD hearts at baseline, with a significant increase in beat-to-beat variability after Iso treatment (Fig 3G and H). Dantrolene suppressed APD alternans after Iso treatment (Fig 3G and H). These data demonstrate that blockade of RyR2 in CIHD hearts prevents arrhythmogenic VERP shortening of the myocardium and reduces the electrical heterogeneity of the myocardium without affecting the conduction velocity, thereby preventing reentrant circuits around the established infarct.

**Figure 3.**
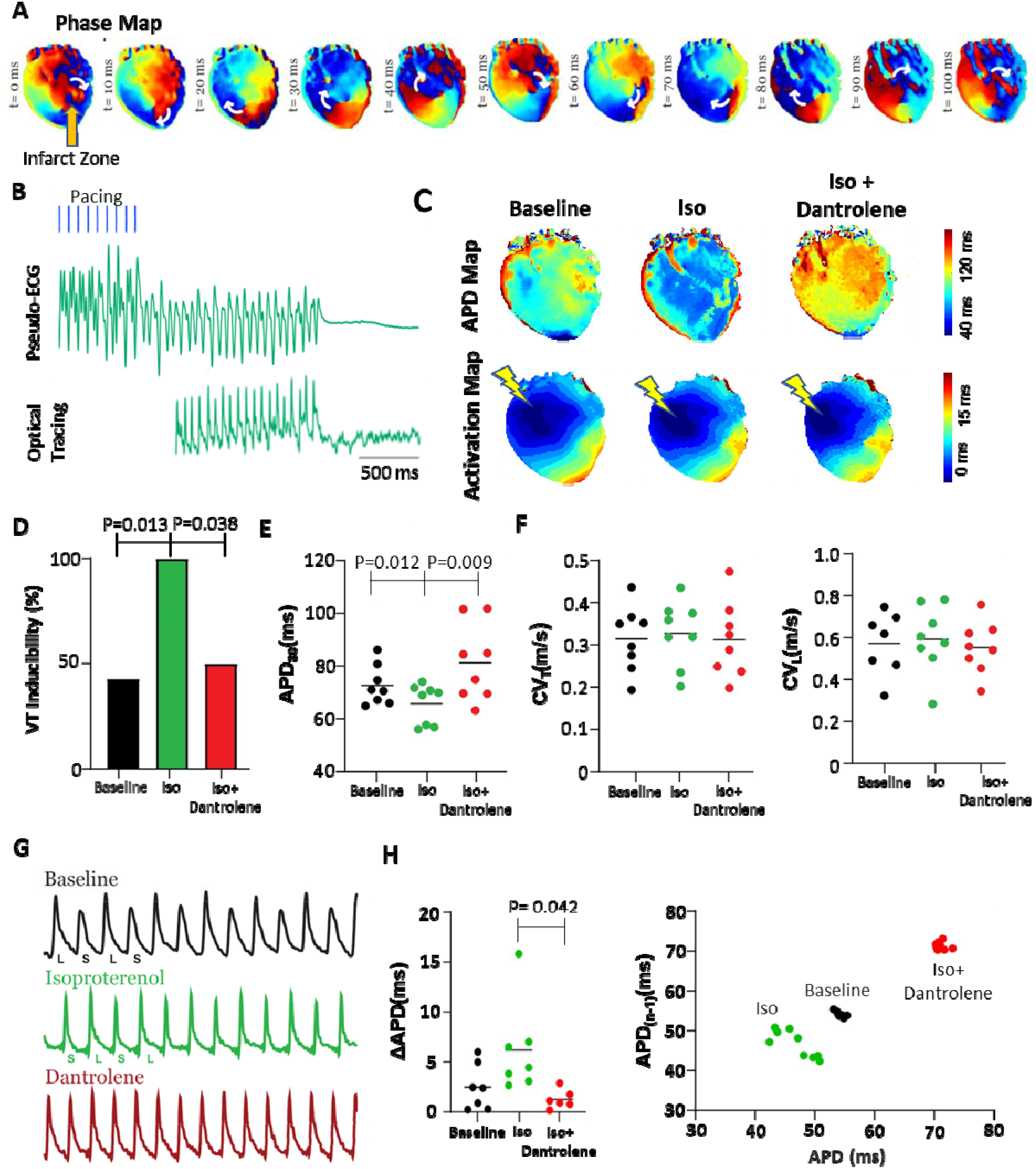
Dantrolene suppresses VT induction in ex vivo CIHD hearts by increasing APD and inhibiting APD alternans. (A) Phase maps illustrating reentrant arrhythmia in an *ex vivo* heart treated with Iso. (B) Volume-conducted ECG and optical (voltage) trace of reentrant ventricular tachycardia (VT) during Iso treatment. (C) APD (top) and activation (bottom) maps during Baseline, Iso and Iso + Dantrolene conditions. (D) VT inducibility, (E) APD80, (F) CVT and CVL during Baseline, Iso and Iso + Dantrolene conditions. (G) Representative optical (voltage) traces demonstrating APD alternans under Baseline and Iso conditions. (H) Summary alternans amplitude during Baseline, Iso and Iso + Dantrolene conditions (left) and Poincaré plot demonstrating increased beat-to-beat variability during Iso treatment (right). P-values were obtained using paired, two-tailed Student’s t-tests with Bonferroni correction for multiple comparisons. [N = 8 mice].

### Acute dantrolene administration suppresses spontaneous SR Ca2+ release from RyR2

Several studies have reported that CIHD renders RyR2 channels hyperactive. To directly assess RyR2 function in our CIHD mouse model, we measured Ca2+ sparks, an indicator of the rate of spontaneous RyR2 openings in cardiomyocytes (Fig 1A). Indeed, Ca+2 sparks frequency (Sham 0.75±0.02 sparks/caff amp vs. CIHD 0.96±0.02 sparks /caff amp, P=2.20×10^−16^; Fig 4B) and Ca2+ leak (Sham: 46.8, 95% CI 0.72-076 vs CIHD 75.7, 95% CI 66.5-84.7, P=6.14×10^−6^; Fig 4C) were significantly increased in ventricular cardiomyocytes isolated from CIHD hearts. Dantrolene administration normalized Ca2+ spark frequency and Ca+2 leak to values observed in cardiomyocytes isolated from sham hearts (Fig 4B-C, Supplemental Fig 2). We next examined Ca2+ handling in intact cardiomyocytes (Fig 4D). After a rapid pacing train with 1 & 3 Hz, isolated myocytes showed increased spontaneous Ca2+ release events, indicative of delayed afterdepolarizations (DADs), which were suppressed by dantrolene (Fig 4D&E). Together, these data showed that dantrolene suppresses spontaneous SR Ca2+ release in CIHD cardiomyocytes.

**Figure 4:**
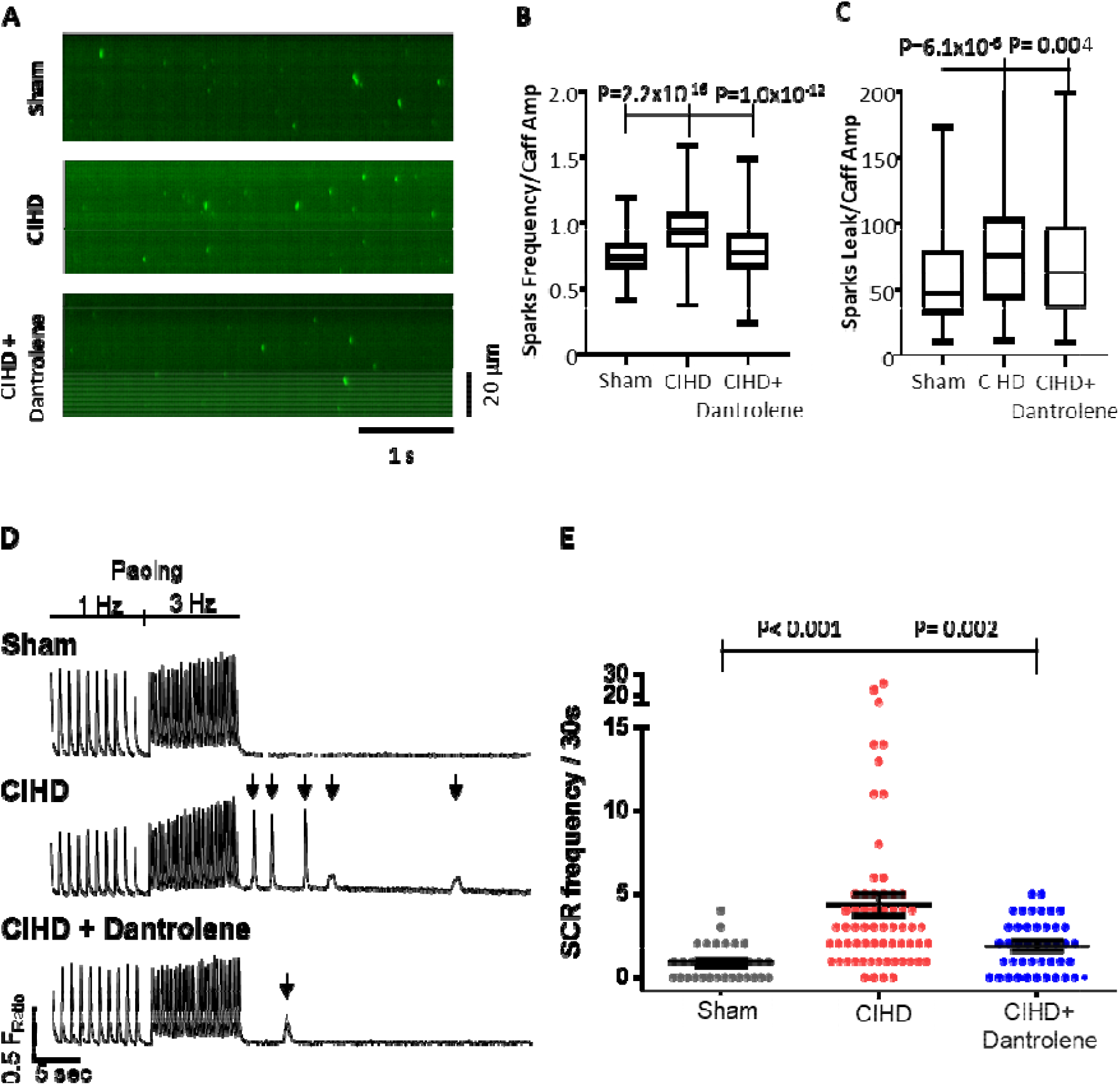
Dantrolene reduces SR Ca2+ leak and spontaneous SR Ca release in CIHD cardiomyocytes. (A) Representative images of Ca Sparks from permeabilized CIHD cardiomyocytes (4 weeks) treated acutely with vehicle or dantrolene. (B) Spark frequency and (C) spark-mediated SR Ca leak in permeabilized cardiomyocytes. [Sham N= 3 mice, n= 121 cells; CIHD N= 3 mice, n= 125 cells; CIHD+Dantrolene N=3 mice, n= 135 cells] (D) Representative Ca transient records from intact cardiomyocytes treated with vehicle or dantrolene. Arrow indicates spontaneous SR Ca release events after the pacing train. (E) Summary data of spontaneous SR Ca release events during a 30s period following the pacing train. P-values obtained using Linear mixed-effect (hierarchical) model with Bonferroni correction. [Sham N=3 mice, n= 33 cells; CIHD N=3 mice, n=64 cells; CIHD+Dantrolene N=3 mice, n= 43 cells]

### Chronic dantrolene treatment suppresses VT risk with minimal adverse effects

Dantrolene has been used clinically for malignant hyperthermia and spasticity but is not without adverse effects, most notably on skeletal muscle weakness, GI symptoms, and liver toxicity. CIHD mice were implanted with subcutaneous osmotic pumps to deliver 20 mg/kg/day of dantrolene over 6 weeks. To test if chronic dantrolene treatment adversely affects CIHD mice, mouse weight and muscle strength were monitored. There was no difference in mortality over the course of the study (Fig 5A). Mice treated with dantrolene show normal growth over the study (Supplemental Fig 2), with mild skeletal muscle weakness noted on EMG (Fig 5B).

**Figure 5:**
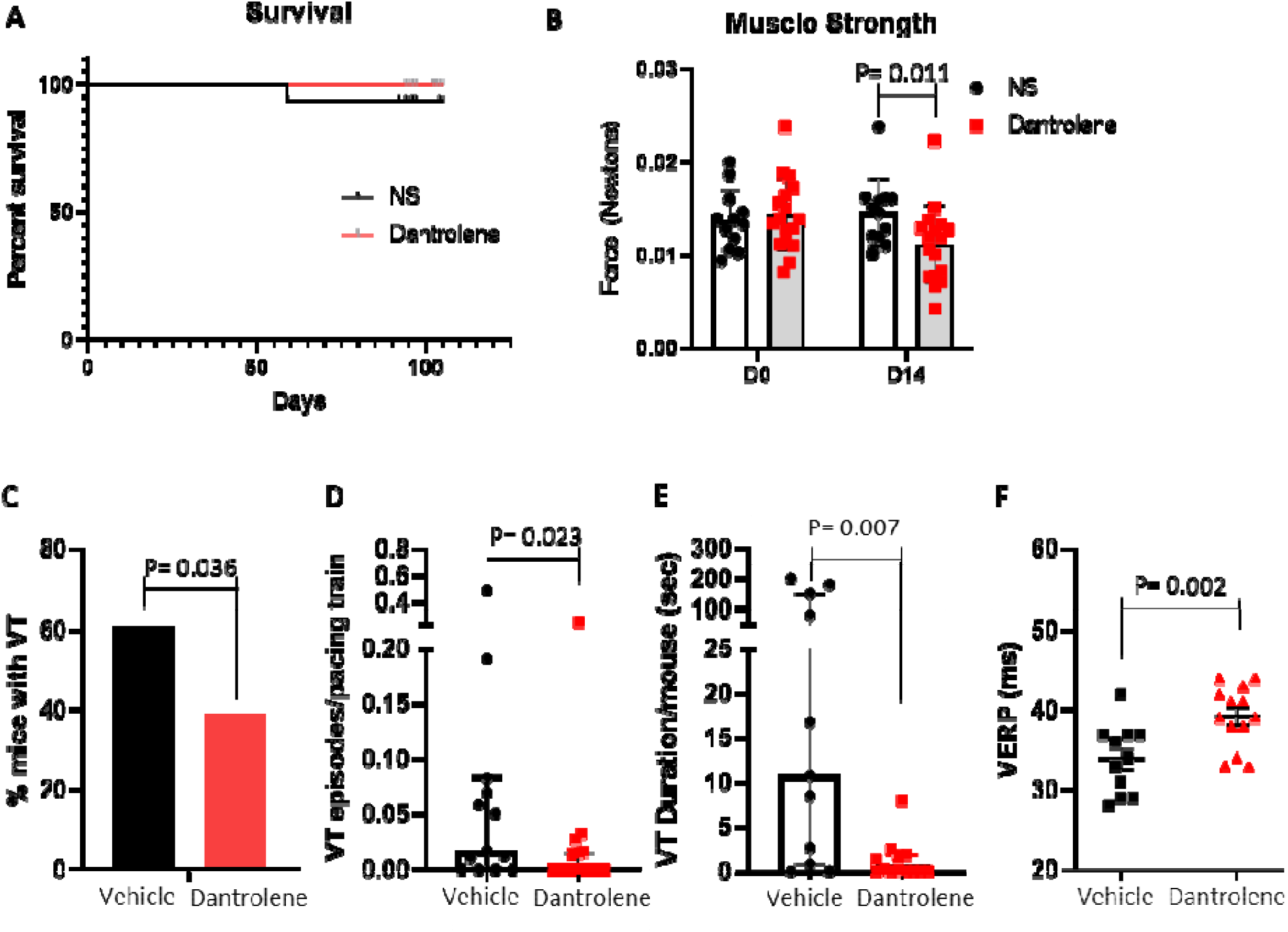
Chronic dantrolene treatment suppresses VT induction with minimal side effects. (A) Survival from time of osmotic pump implantation (4 weeks post-MI). (B) Mice showed a significant decrease in peripheral skeletal muscle strength at 2 weeks of dantrolene treatment. Effect of 6-week dantrolene treatment on (C) VT inducibility, (D) VT episodes/pacing train, (E) VT duration and (F) ventricular effective refractory period (VERP). P values were obtained using the Wilcoxon signed-rank test. [Vehicle n=13, Dantrolene n=17]

Dantrolene treatment suppressed VT induction by 22.2% (61.1% vehicle vs 38.9% dantrolene; P=0.036 Fischer Exact Test) (Fig. 5C). Dantrolene also significantly reduced the duration and frequency of VT episodes (Fig 5D). As with acute treatment, chronic dantrolene normalized the VERP at 6 weeks (Vehicle 33.9±1.3 ms vs. Dantrolene 39.2±1.1 ms; P=0.032) (Fig 5D). These data show that chronic dantrolene treatment is as effective as an acute treatment for preventing VT in CIHD with minimal long-term adverse effects.

### Chronic dantrolene treatment prevents progressive LV dysfunction and peri-infarct fibrosis in CIHD

LV remodeling after MI through infarct expansion in the border zone, interstitial fibrosis, and chamber dilation provides a substrate for reentry ventricular arrhythmias as well as reducing cardiac function and leading to heart failure. To test if dantrolene can prevent progressive remodeling late after MI, cardiac function, fibrosis, biomarkers for wall stress, and hypertrophy were assessed after 6 weeks of dantrolene treatment.

Although it did not change the size of the infarct scar, dantrolene treatment largely prevented the development of interstitial fibrosis in the peri-infarct zone (Fig 6B). Expression of genes responsible for cardiac fibrosis (Col1a1, Col1a3, Postn), heart failure biomarkers of increased wall stress (Nppa, Nppb), and hypertrophy (Myh7/Myh6 ratio) continued to be upregulated in the infarct border zone at 10 weeks post infarction. However, these were significantly reduced after 6 weeks of treatment with dantrolene (Fig 6C, Supplemental Table 2). Fibrosis was not significantly increased in the remote region. Heart failure (Nppa, Nppb) and hypertrophy (Myh7/Myh6 ratio) biomarkers were also significantly increased in the remote region after infarction, but the expression was less than in the infarct border zone.

**Figure 6:**
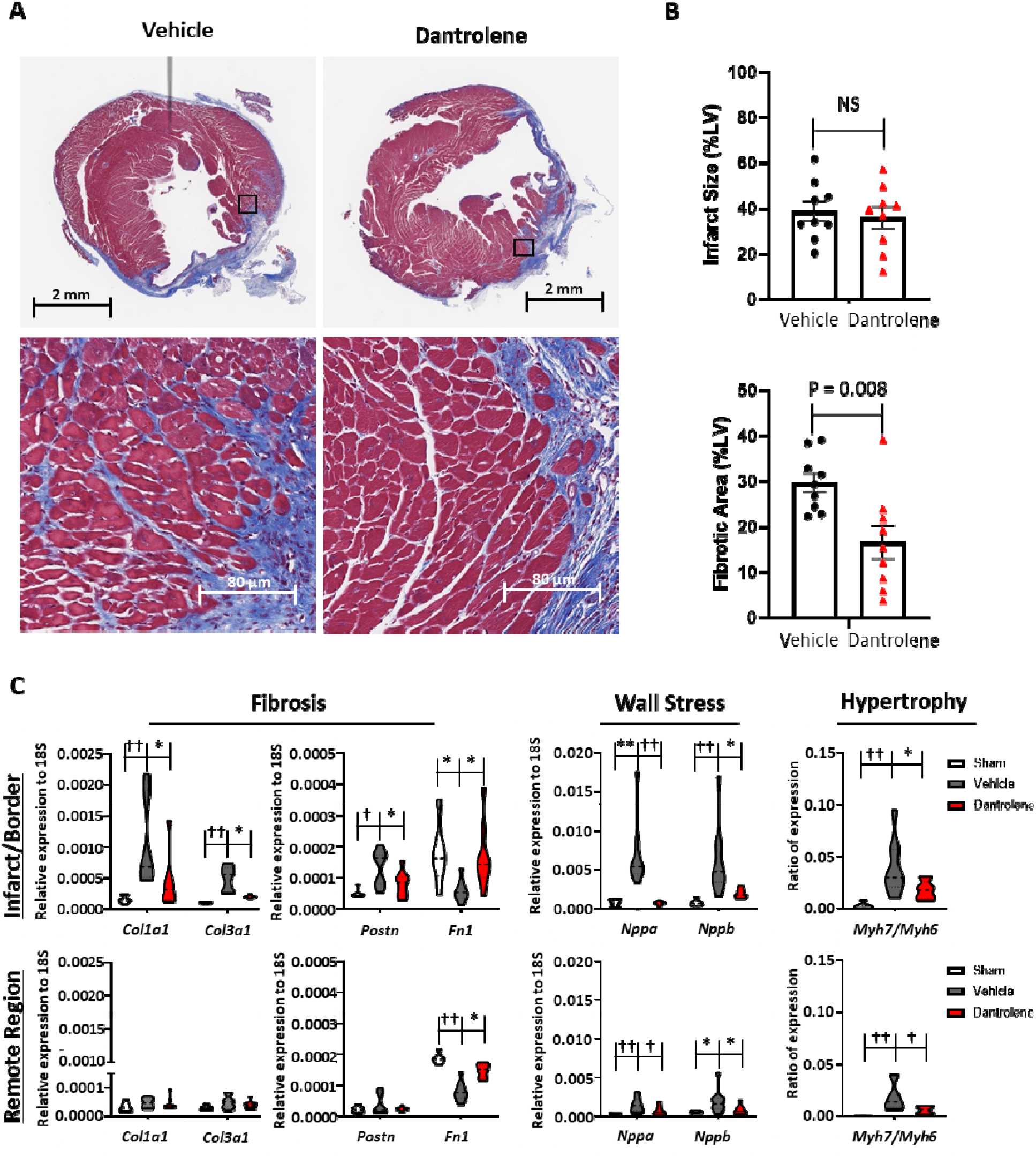
Chronic dantrolene reduces fibrosis and LV remodeling in ischemic cardiomyopathy. (A) Representative mid-LV Masson’s Trichrome staining from CIHD mice after 6 weeks treatment with vehicle or dantrolene. (B) Quantification of infarct scar size and fibrosis. P-value obtained using Mann-Whitney test. [Vehicle n=9, Dantrolene n=9] (C) Dantrolene reduced gene expression of fibrotic, wall stress and hypertrophy markers in infarct/border zone after 6 weeks of treatment. P values were obtained using Kruskal-Wallis test with Dunn’s post-test [* P<0.05, ** P<0.01,† P<.005, †† P<0.001; Sham = 6, Vehicle n=9, Dantrolene n=9]

Concordant with the changes observed in the border zone, these markers were reduced in the remote region with 6-week dantrolene treatment. Notably, dantrolene also normalized the reduced fibronectin (Fn1) expression in the infarct border zone and remote region 10 weeks (Fig. 6). These data show that treatment with dantrolene reduces progressive fibrosis and negative remodeling in both the border zone and remote region in CIHD mice.

Chronic dantrolene treatment significantly improved LV contractile function compared to vehicle, as evidenced by an increased fractional shortening (Fig 7A; Supplemental Table 3) and mean velocity fiber shortening (Fig 7B; Supplemental Table 3) with 6-week dantrolene treatment. Consequently, dantrolene prevented the progression of systolic dysfunction (Fig 7C; Supplemental Table 3) and reduction in LV ejection fraction from 4 to 10 weeks post-infarction (Fig 7D; Supplemental Table 3). These data demonstrate the therapeutic efficacy of dantrolene for preventing progressive pathological remodeling and improve cardiac function even after heart failure has been established.

**Figure 7:**
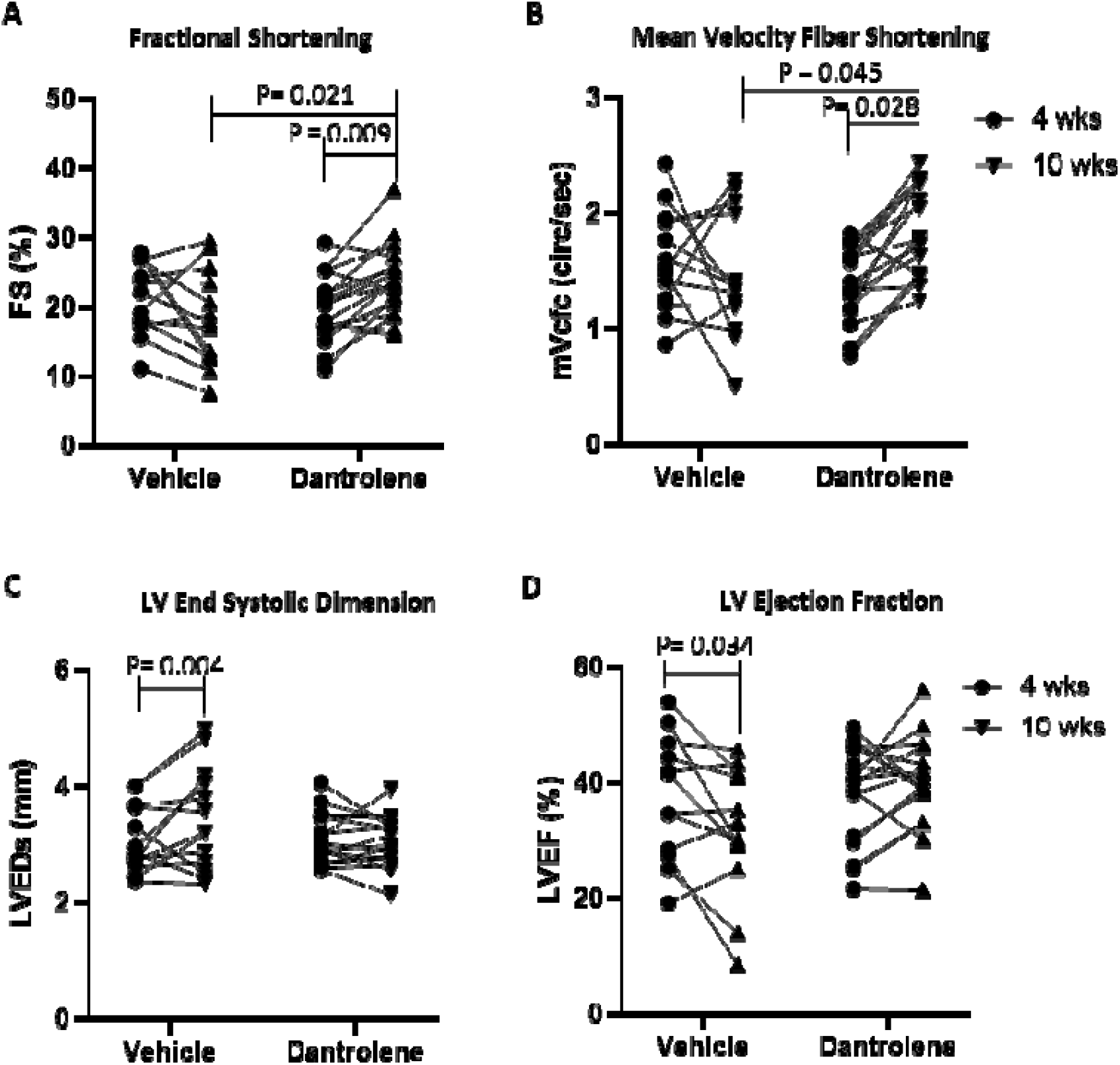
Chronic Dantrolene treatment improves cardiac function in the CIHD model. Dantrolene improved (A) fractional shortening and (B) contractility after 6 weeks of treatment. Dantrolene prevented progressive (C) systolic dysfunction and (D) LVEF decline after 6 weeks of treatment. P-value obtained using 2way ANOVA mixed-effect model with Bonferroni post-test. [Vehicle n=13, Dantrolene n=17]

## Discussion

Persistent LV systolic dysfunction 40 days after MI is associated with increased mortality from SCD due to ventricular arrhythmias. While ICDs and current heart failure guideline directed medical therapy have improved mortality, a third of patients continue to have progressive LV dysfunction, which increases the risk of ventricular arrhythmias and can ultimately lead to the need for cardiac transplant and increased risk of ventricular arrhythmias.^31^ Therapy with ICD placement, anti-arrhythmic drugs and/or catheter ablation reduces the risk of ventricular arrhythmias. However, current anti-arrhythmic drug therapy for VT in structural heart disease is primarily limited to Class III drugs and beta-blockers, which have issues with drug toxicities, intolerance and/or limited effectiveness. The use of dantrolene to suppress RyR2 hyperactivity in the failing heart represents a new therapeutic target to prevent VT and improve cardiac function in CIHD.

Cardiac remodeling after an ischemic injury has been shown to induce both areas of fixed anatomical block in the setting of fibrosis and functional block by slowing conduction or changing the refractoriness of the surviving myocardium.^32^ The mechanisms of ventricular arrhythmias VT in CIHD include a combination of triggered activity beats due to diastolic SR Ca2+ release, resulting in DADs, and substrate heterogeneous refractoriness due to interstitial fibrosis that allows for reentry. Diminished cell-to-cell coupling associated with fibrosis and abnormal Ca2+ can contribute to slow conduction and provide reentry substrate. In addition, functional block and APD alternans due to abnormal Ca2+ handling drives T-wave alternans and is associated with SCD.^33, 34^ Continued diastolic SR Ca2+ leak also leads to poor contractile reserve in the surviving myocardium, worsening cardiac function. The most crucial finding in this study is that inhibition of RyR2 calcium leak not only suppressed inducible VT but also reduced cardiac fibrosis and improved cardiac function. The prevention of further negative remodeling after MI is even more significant considering treatment with dantrolene was not started until 4 weeks after coronary ligation, when the infarct scar was already established. This demonstrates the utility of dantrolene and RyR2 inhibition in the heart to prevent VT and improve systolic function with the surviving myocardium late after injury.

### Effect of Dantrolene on Cardiac Electrophysiology in CIHD

It has been recognized that altered Ca2+ cycling produces Ca2+ alternans, which contribute to the formation of functional reentry and arrhythmia by inducing the formation of functional reentry and arrhythmia by inducing dispersion of excitability or refractoriness, and is associated with SCD. ^35, 36^ Transmural heterogeneity of APD in CIHD rabbits has shown a correlation between the beat-to-beat cycle length at which alternans occurs and a reduction in the VF threshold.^37, 38^ Targeting functional reentry and substrate heterogeneity in infarcted tissue is key to preventing ventricular arrhythmias and SCD. Class Ic anti-arrhythmic drugs can induce electrical alternans^39^ and ventricular arrhythmias^40^ in ischemic hearts, and are contraindicated in structural heart disease.^41, 42^ One of the most striking results of this study is that dantrolene normalizes APD and VERP in CIHD and suppresses APD alternans and substrate heterogeneity in the infarct border zone without affecting conduction time.

Several animal models have demonstrated the utility of suppressing RyR2 diastolic Ca2+ leak in preventing reentry mechanisms in AF, VT and VF.^12, 22, 24^ The proposed mechanism of dantrolene is to stabilize the tertiary structure, which prevents Ca2+ leak. Consistent with prior data, dantrolene treatment suppressed spontaneous SR Ca2+ release and Ca2+ sparks in the CIHD model. Notably, both acute and chronic dantrolene treatment reduced VT induction by normalizing the APD and VERP *in vivo* with minimal adverse effects. The data here provide proof of concept for another therapeutic option to patients with breakthrough ventricular arrhythmias where other anti-arrhythmic drugs have failed or are not tolerated.

### Effect of Dantrolene on Cardiac Function in CIHD

RyR2 hyperactivity leading to diastolic Ca2+ leak has been documented in isolated myocytes from animal models and patients with systolic heart failure.^43^ Increased Ca2+ leak reduces SR Ca2+ stores, which subsequently diminishes Ca2+ release in systole, leading to reduced myocyte contraction. Thus, stabilizing RyR2 and preventing diastolic Ca2+ leak could improve cardiac function in HF. Our data show suppression of RyR2 hyperactivity by dantrolene reduces pathological cardiac remodeling and improves LV function.

Few studies have evaluated the effect of RyR2 inhibition on cardiac function and remodeling in CIHD. Isolated cardiac strips from patients with idiopathic dilated cardiomyopathy have improved force generation with dantrolene treatment in response to isoproterenol.^20^ In dogs with RV pacing-induced cardiomyopathy, long-term treatment with dantrolene blunted chamber dilation and improved LV systolic function over 4 weeks.^21^ Recently, in a rat model of MI, treatment with dantrolene at the time of MI improved LV function, reduced atrial fibrosis and atrial fibrillation, although infarct size and LV fibrosis were not reported in this study.^16^ In contrast to these animal models where dantrolene was administered at the time of injury, in our model of CIHD, mice have an established infarct scar prior to dantrolene treatment, which is more clinically relevant to patients with ischemic cardiomyopathy.

In our model, long-term dantrolene treatment reduced border zone fibrosis and myocyte hypertrophy while improving markers of wall stress and cardiac function. A possible mechanism here is that improved Ca2+ handling in the surviving cardiomyocytes have restores contractile function leading to reduced wall stress, as seen with reduced ANP and BNP expression. This would attenuate the pathological remodeling driven by cytokine and neurohormonal activation.^13^ Interestingly, the expression of fibronectin was normalized in the infarct border zone and remote region late after infarction. While fibronectin enhances collagen type I polymerization and fibrosis early after infarction^44^, Fn1-KO mice show progressive LV dysfunction at later time points after MI.^45^ Overall, these results support targeting RyR2 hyperactivity late after ischemia to suppress progressive negative remodeling of the heart by reducing ongoing fibrosis, wall stress, and hypertrophy in the infarct border zone to improve cardiac function in ischemic heart failure.

In conclusion, our data provide proof of concept for targeting RyR2 hyperactivity and diastolic Ca2+ leak to suppress ventricular arrhythmias, reduce cardiac fibrosis and improve cardiac function in CIHD.

## Non-standard Abbreviations and Acronyms

AF: Atrial Fibrillation
APD: Action Potential Duration
CIHD: Chronic Ischemic Heart Disease
CPVT: catecholaminergic polymorphic ventricular tachycardia
DAD: delayed afterdepolarization
EMG: Electromyogram
FS: Fractional shortening
ICD: implantable cardioverter-defibrillator
PSAX: parasternal short-axis
PSLX: parasternal long-axis
LV: Left ventricle/ventricular
LVEDd: Left ventricular end-diastolic dimension
LVESd: Left ventricular end-systolic dimension
LVEF: Left ventricular ejection fraction
MI: Myocardial Infarction
mVcfc: mean velocity circumferential fiber shortening
PES: Programmed electrical stimulation
RV: Right ventricle/ventricular
RyR2: Ryanodine Receptor 2
SR: Sarcoplasmic reticulum
SCD: Sudden cardiac death
VT: Ventricular Tachycardia
VF: Ventricular Fibrillation

## Acknowledgments

We acknowledge the Translational Pathology Shared Resource supported by NCI/NIH Cancer Center Support Grant P30CA068485 and Shared Instrumentation Grant S10 OD023475-01A1 for the Leica Bond RX.

