## Supplemental Methods for "RyR2 inhibition with dantrolene is antiarrhythmic, antifibrotic, and improves cardiac function in chronic ischemic heart disease"

**Supplemental Material**

**Methods**:

**Mouse CIHD model:** All studies were approved by the Vanderbilt Animal Care and Use Committee of Vanderbilt University, USA (Protocol M1900081-00) and performed in accordance with NIH guidelines. To induce CIHD, 10-12 week old male and female C57BL/6J mice underwent complete ligation of the left coronary artery as previously described.^25^ Mice were allowed to recover for 4 weeks before inclusion into the study. The inclusion criteria for CIHD mice were 1) Fractional shortening < 35%, 2) Ejection fraction <50%, and 3) mean velocity fiber shortening <2.5 circ/sec. A total of 39 male and 54 female mice were included in the study. All data analysis was performed in a blinded fashion regarding the treatment group.

For chronic treatment of dantrolene, osmotic mini-pump (#2006, Alzet) were implanted in CIHD mice 4 weeks post coronary ligation as previously described. Dantrolene sodium suspension (Ryanodex, Eagle Pharmaceuticals) was diluted in 0.9% normal saline to deliver 20 mg/kg/day of dantrolene over 6 weeks.

**Mouse Echocardiography:** Infarcted mice underwent longitudinal echocardiography to assess cardiac function at 4 and 10 weeks post-MI. Mice are anesthetized with 1.5% isoflurane and placed on a heated stage for the procedure. B-mode images were taken in the parasternal long-axis (PSLX), 2-chamber, and 4-chamber views. B-mode and M-mode were taken at the base parasternal short axis (PSAX) just below the tip of the left atrial with en face view of the mitral valve tips, mid ventricle PSAX at the level above the insertions of papillary muscles, and apex PSAX below the level of the papillary muscles. Color flow and pulse Doppler were taken at the mitral valve in the 4-chamber view. Fractional shortening (FS), left ventricular end-diastolic dimension (LVEDd) and left ventricular end-systolic dimension (LVESd) in CIHD mice were calculated from the average of the PSAX M-mode at the base, mid ventricle and apex. The left ventricular ejection fraction was measured using Simpson’s method in the PSLX and PSAX B-mode views. Contractility was calculated by mean velocity circumferential fiber shortening (mVcfc) by the equation mVcfc = (LVEDd-LVESd / LVEDd) x (√(60/HR)/LV ejection time). All images were analyzed using Vevo Lab software (FUJIFILM VisualSonics, Toronto, ON)

**Mouse transesophageal programmed electrical stimulation (PES):** Sham and CIHD mice were anesthetized with inhaled isoflurane (3% for induction, 2-2.5% for maintenance) while breathing spontaneously and placed in the supine position on a heating pad. Surface ECG was recorded continuously using AD Instruments amplifiers and LabChart 8 software. An octopolar 2F electrode catheter (CIB’ER MOUSE™; NuMED, Inc) was placed in the esophagus via the mouth, guided by electrogram tracings to verify position. Unipolar pacing was performed using a programmable stimulator with 6 mA of pacing amplitude and 3 ms of pulse width for all studies. PES consisted of pacing with a train of 15 beats (10 Hz, S1), followed by a single extra stimulus (S2) to determine the ventricular effective refractory period (VERP). VT induction was then performed using 3 extra stimuli (S2-S4) following each pacing train to induce VT. Pacing trains were repeated every 5 seconds for a total 30 sets per animal. Inducible VT was defined as a ventricular arrhythmia consisting of at least three consecutive beats following each pacing train. At the end of the procedure, the esophageal catheter and subcutaneous needles are removed, and isoflurane was discontinued. The animal was placed in a cage with food and water and observed until recovery. Dantrolene (30 mg/kg, intraperitoneal injection) using Ryanodex (Eagle Pharmaceuticals, Inc., NJ) or normal saline was administered to mice 30 minutes prior to the study. Isoproterenol (1.5 mg/kg, intraperitoneal injection) was administered after capturing ventricular pacing.

**Isolation of ventricular myocytes:** Ventricular myocytes from CIHD or Sham C57BL/6J wild-type mice were isolated from the left ventricle by collagenase/protease digestion as previously described.^26^ Cells were then washed twice by gravity sedimentation for 20 minutes at room temperature in standard Tyrode solution with 0.2 mM CaCl2. The final suspension contained 0.6 mM Ca2+ and the cells were immediately used for Ca2+ sparks or intact intracellular Ca2+ measurements.

**Mouse electromyography/kinesiology:** Mice undergoing chronic drug treatment with osmotic pumps were monitored for weakness by hindlimb electromyography. Mice were anesthetized with inhaled isoflurane (2%) while breathing spontaneously and placed prone on a heating pad. Needle electrodes were placed 1 cm apart in the sciatic notch and were stimulated with 15 pulses of 5 ms, 10 mA to induce repeated contraction of the gastrocnemius muscle. An electromyogram (EMG) was recorded from needle electrodes in the gastrocnemius muscle. Maximum muscle strength was measured with a force transducer (AD Instruments) placed over the paw. Stimulated pulses and recordings were acquired using PowerLab 26T (AD Insturments) and analyzed using LabChart Pro 4 software (AD Insturments).

**Histology of mouse hearts:** Hearts from CIHD mice were harvested at 10 weeks post coronary ligation. Mice were anesthetized with 2.5% isoflurane and were injected with ice-cold cardioplegic solution (110 mM NaCl, 10 mM NaHCO3, 16 mM KCl, 16 mM MgCl2, and 1.2 mM CaCl2, 1000 units/mL heparin) by intraventricular injection to arrest the heart in diastole. A 1 mm mid-ventricular slice distal to the ligation site through the infarct zone was taken and frozen in liquid nitrogen for RNA isolation. Hearts were fixed in Tris-buffered 4% paraformaldehyde solution overnight, paraffin-embedded for sectioning, and stained by H&E and Mason’s Trichrome methods. 3 noncontiguous sections from each animal were analyzed for infarct size and fibrosis area. Infarct size was determined using ImageJ to calculate the percentage of the circumferential length of the scar to the total left ventricular circumference in mid-ventricular axial H&E sections. Fibrosis was determined using ImageJ by identifying blue staining on Mason’s Trichrome section and calculated as the percentage of the left ventricle total area.

**Quantitative Reverse Transcriptase PCR:** Frozen heart sections from CIHD mice were harvested as above. The RV was removed and the LV was separated into the infarct border zone and remote region. RNA was isolated using TriZol Reagent (Thermo Fisher) per the manufacturer’s protocol. cDNA was generated using Invitrogen SuperScript IV VILO (Thermo Fisher) per the manufacturer’s protocol. Gene expression for markers of fibrosis (Collagen type I [Col1a1: Mm00801666_g1] ,Collagen type III [Col1a3: Mm00802300_m1], Periostin [Postn: Mm01284919_m1],fibronectin [Fn1: Mm01256744_m1]), Hypertrophy (α-myosin heavy chain [Myh6: Mm00440359_m1], β-myosin heavy chain [Myh7: Mm00600555_m1]), and heart failure biomarkers (ANP [Nppa: Mm01255747_g1], BNP [Nppb: Mm01255770_g1]) were measured using TaqMan Gene Expression Assay and measured using an Applied Biosystems QuantStudio 3 (Thermo Fisher). Each reaction was done on 100 ng of cDNA and the 2^-ΔCT^ method was used to calculate the relative expression to 18S RNA [Hs99999901_s1].

**Optical mapping of isolated mouse hearts:** CIHD mice were anesthetized by isoflurane 4 weeks after coronary ligation. Hearts were isolated after thoracotomy and Langendorff perfused with a modified Tyrode's solution (130 mM NaCl, 24 mM NaHCO3, 1.2 mM NaH2PO4, 4 mM KCl, 1 mM MgCl2, 5.6 mM Glucose and 1.8 mM CaCl2, pH 7.4 at 37⁰C). Cardiac motion was arrested during optical mapping using blebbistatin (15 μM). Hearts were paced using a bipolar platinum pacing wire placed on the anterior surface of the heart, at the center of the field of view, using pulses at 1.5x threshold of stimulation and 2 ms duration. Restitution properties were measured by pacing at multiple basic cycle lengths (BCL) from 200 - 60 ms. After a 15 min equilibration period, hearts were stained with di-4-ANEPPS, a voltage sensitive dye (37.5 μg/ml in Tyrode solution). After a 5 min washout period, the dye was excited using light at 510±5 nm wavelength. Emitted fluorescence was filtered using a 610±20nm bandpass filter and recorded using a CMOS camera (MiCam05, SciMedia). Optical recordings were obtained at baseline, Isoproterenol (250 nM) treatment and Iso + Dantrolene (10 μM) treatment. Optical signals were analyzed using a custom Matlab program (Rhythm).

**Ca2+ transient measurements in intact cardiomyocytes:** Cardiomyocytes were loaded with Fura-2 acetoxymethyl ester (Fura-2 AM; Invitrogen) as described previously.^27^ Briefly, isolated single ventricular myocytes were incubated with 2 µM Fura-2 AM for 6 minutes, washed 2 times for 10 minutes each with normal Tyrode (NT) solution containing 2.5 µM probenecid and 1.2 mM CaCl2. The composition of NT solution was: 134 mM NaCl, 5.4 mM KCl, 10 mM glucose, 1 mM MgCl2, and 10 mM HEPES (pH adjusted to 7.4 with NaOH). After Fura-2 loading, experiments were conducted in NT solution containing 1 µM isoproterenol and 2 mM CaCl2. Fura-2 AM-loaded myocytes were pre-incubated for 1 hour with vehicle or 1 µM dantrolene. Myocytes were then electrically paced at 1 and 3 Hz field stimulation for 10 seconds each, followed by no electrical stimulation for 30 seconds. At room temperature, the intracellular Ca2+ ratio was recorded using a dual-beam excitation fluorescence photometry setup (IonOptix Corp.). Spontaneous Ca2+ release events were quantified during the 30 seconds following cessation of the pacing train.

**Ca2+ spark measurements in permeabilized cardiomyocytes**: Ca2+ sparks were measured in isolated adult myocytes from CIHD mice by spinning-disc confocal microscopy as previously described.^28^ Since RyR2 activity is regulated by both cytosolic and intra-SR Ca2+, cytosolic Ca2+ was clamped to 54 nM by cell membrane permeabilization with saponin and Ca2+ sparks were measured using Fluo-4. Cells were treated with vehicle (0.1% DMSO) or 10 μM dantrolene for 30 minutes prior to imaging. Calcium sparks were recorded at room temperature on an Olympus spinning disk confocal microscope equipped with 488 nm diode laser and filters. Analysis of spark frequency was performed using the SparkMaster plugin for ImageJ and all data were normalized to SR Ca2+ content for the experimental day. Statistical comparisons were made using a linear mixed-effects (hierarchical) model provided by Sikkel et al., clustered by mouse to account for random effects between isolations and calculate Bonferroni-adjusted p values.

**Statistics**: Statistical analyses were performed using Prism v9.0.2 (GraphPad Software, Inc.). Statistical tests that were used are reported in the statistical summary table (Supplemental Figure 1) and are also reported in the figure legends. Data were tested for normality using the Shapiro-Wilk normality test and p-values for each set of data are included in the statistical summary table. For normally distributed data (VERP, APD_80_, Infarct Size, Fibrosis area, FS, mcVcf, LVESD, LVEF), mean and standard deviation are provided. ANOVA with Bonferroni correction was used for multiple comparisons to assess for interaction between variables. For non-normally distributed data (VT episodes/train, VT duration, Ca2+ sparks, and Spontaneous Ca2+ release), the mean and 95% confidence intervals are provided. Linear regression models were used to assess the interaction between multiple variables and multiple comparisons when the data were non-normally distributed. For Ca2+ sparks analysis, hierarchical clustering was performed by animal in R using the script provided by Sikkel et al. Briefly, a linear mixed effects (hierarchical) model (lmer) was used to calculate Bonferroni-adjusted p values clustered at the animal level to account for random effects between isolations. The script also calculates goodness of fit for both the non-hierarchical and hierarchical models and generates a p value to determine whether the hierarchical model is a significant improvement over the non-hierarchical equivalent. This was used as a justification for using the hierarchical model and is included in the statistical summary table. A corrected p-value of 0.05 was used as the threshold to reject the null hypothesis. Raw and corrected p-values are available in the statistical summary table. Corrections for multiple tests were only made within-tests. The number of corrections made is included in the statistical summary table as part of the supplemental data. The p-values reported in the manuscript text and figures are the p-values corrected for multiple comparisons.

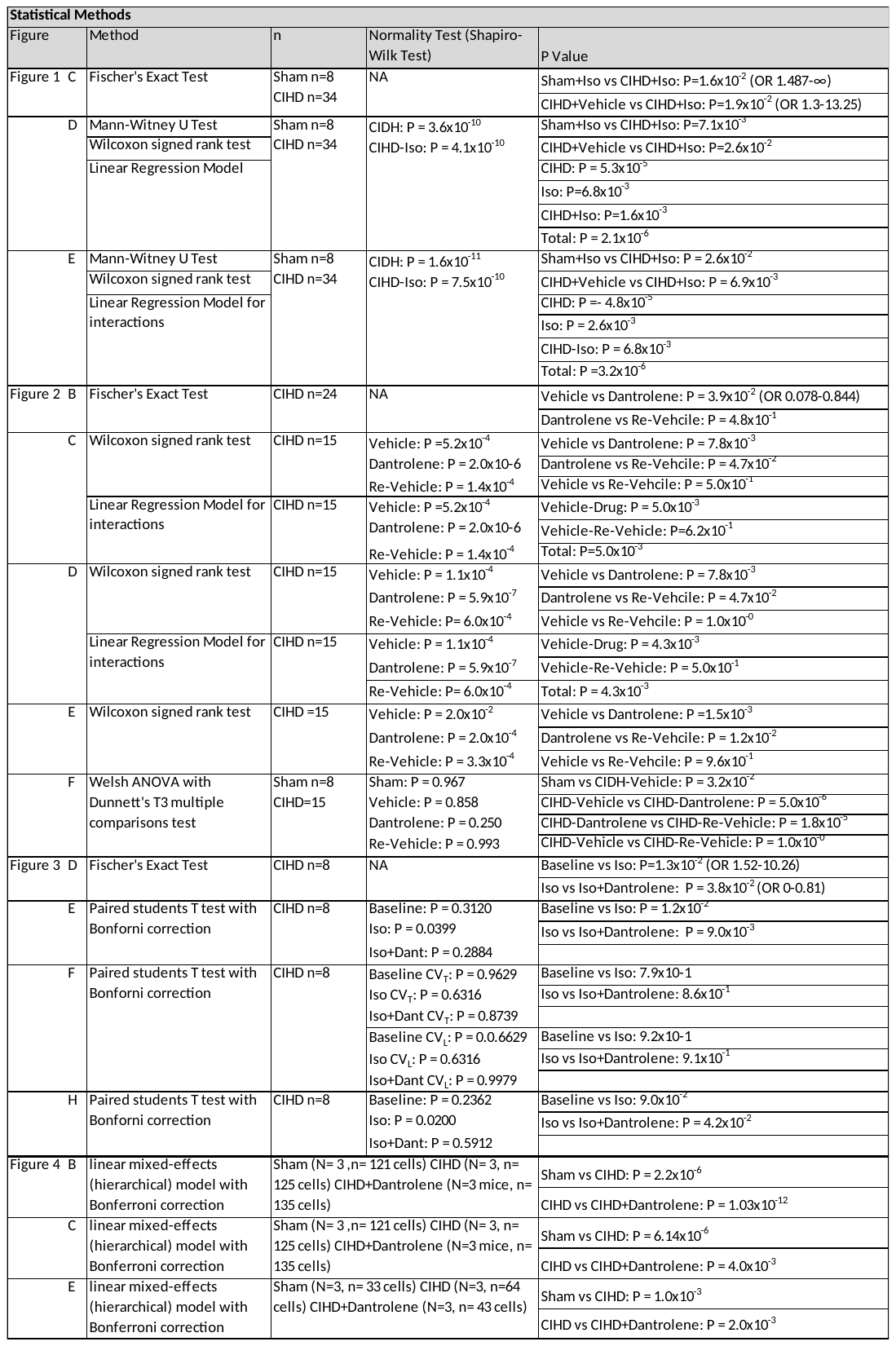

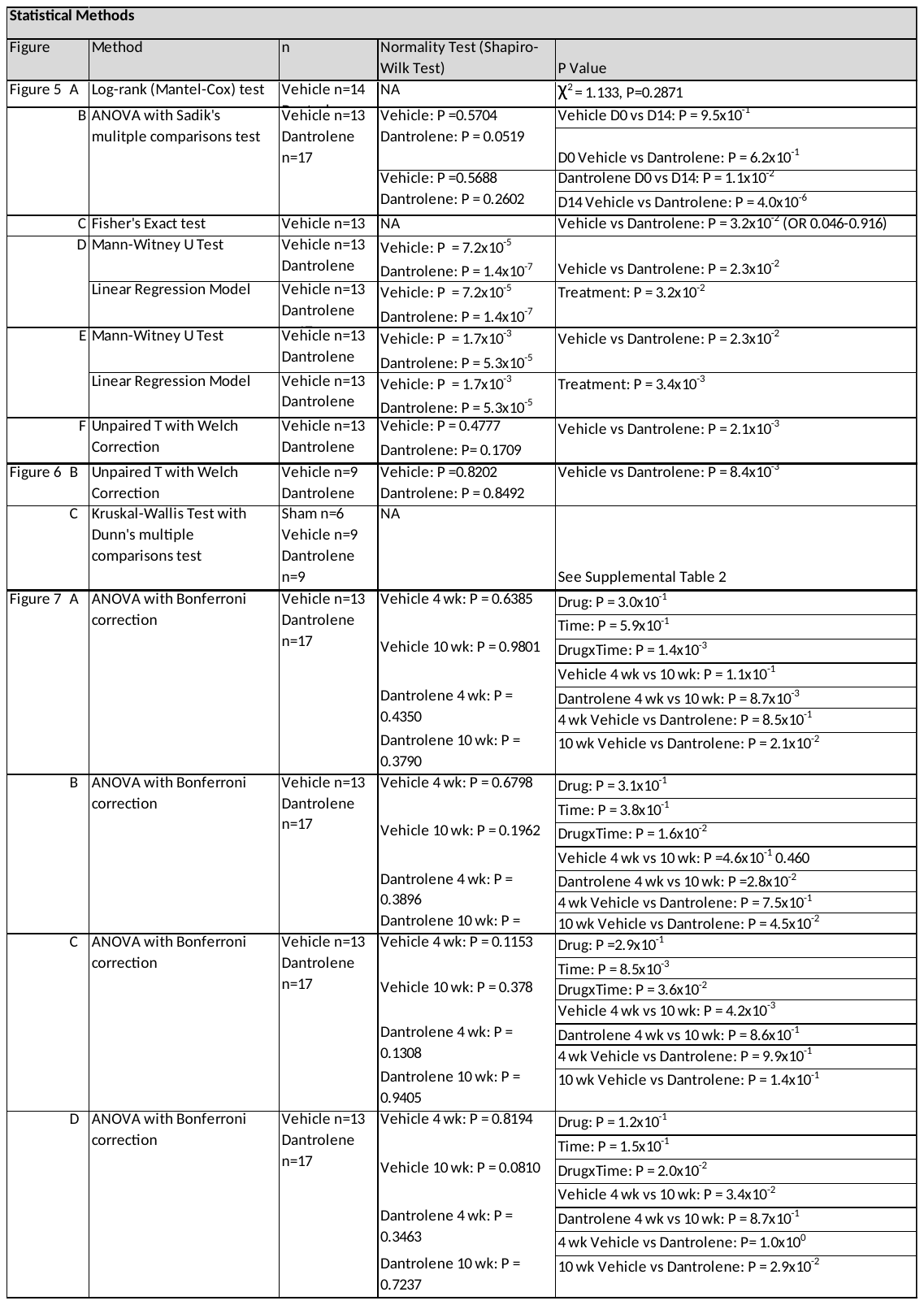

**Sup Table 1: Statistical methods used for analysis by figure**

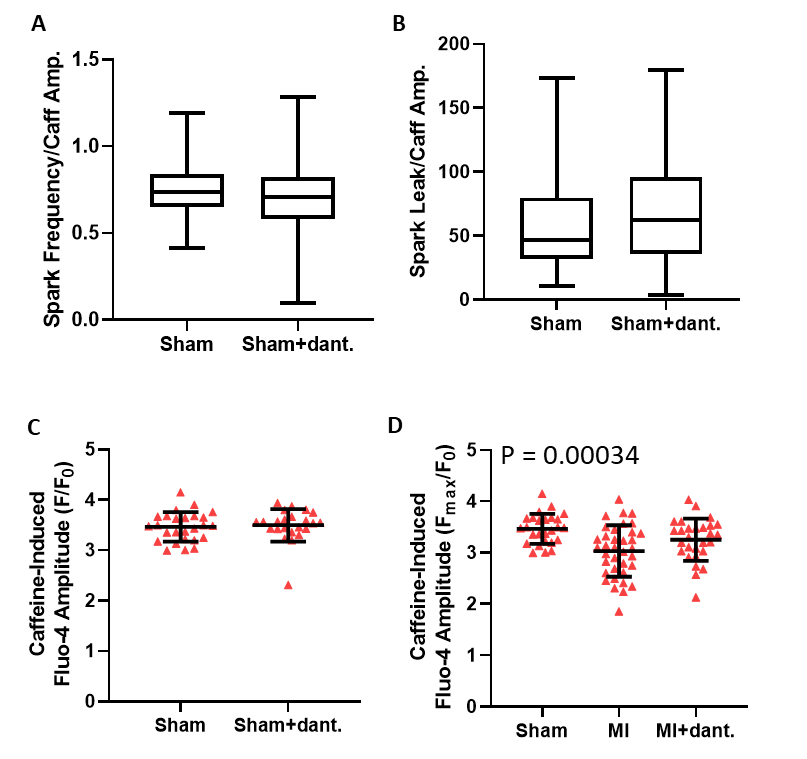

**Sup Figure 1: Dantrolene does not affect Ca+2 sparks in normal cardiomyocytes after cardiac infarction.** (A) Spark frequency and (B) spark-mediated SR Ca leak in permeabilized cardiomyocytes from sham animals with and without dantrolene treatement. [Sham N= 3 mice, n= 121 cells; [=3 mice, n= 33 cells]. Caffeine-induced Ca2+ amplitude in (C) Sham or (D) CIHD mice with and without dantrolene. [Sham N=3 mice, n= 33 cells; CIHD N=3 mice, n=64 cells; CIHD+Dantrolene N=3 mice, n= 43 cells]

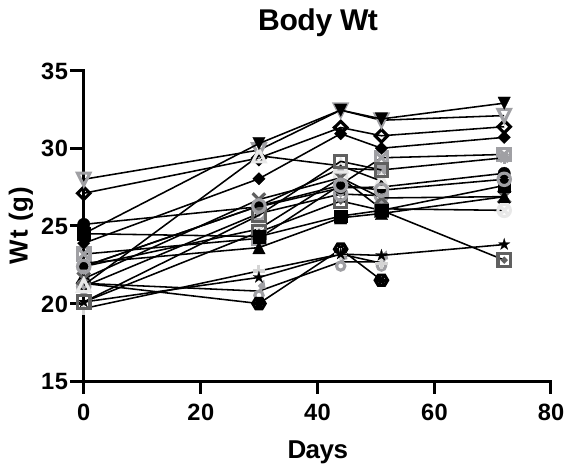

**Sup Figure 2: Chronic dantrolene treatment did not affect normal mouse growth or feeding.** Mouse weight from time of osmotic pump implantation to sacrifice data (6 weeks).

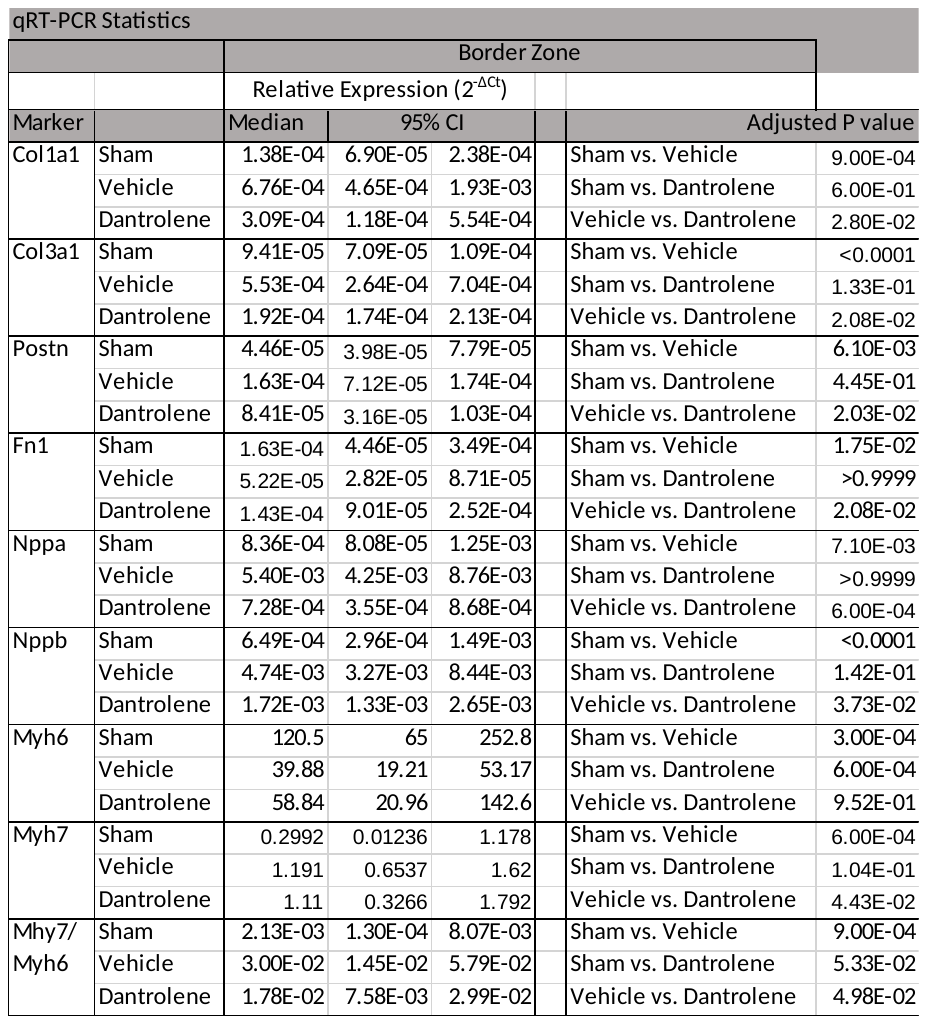

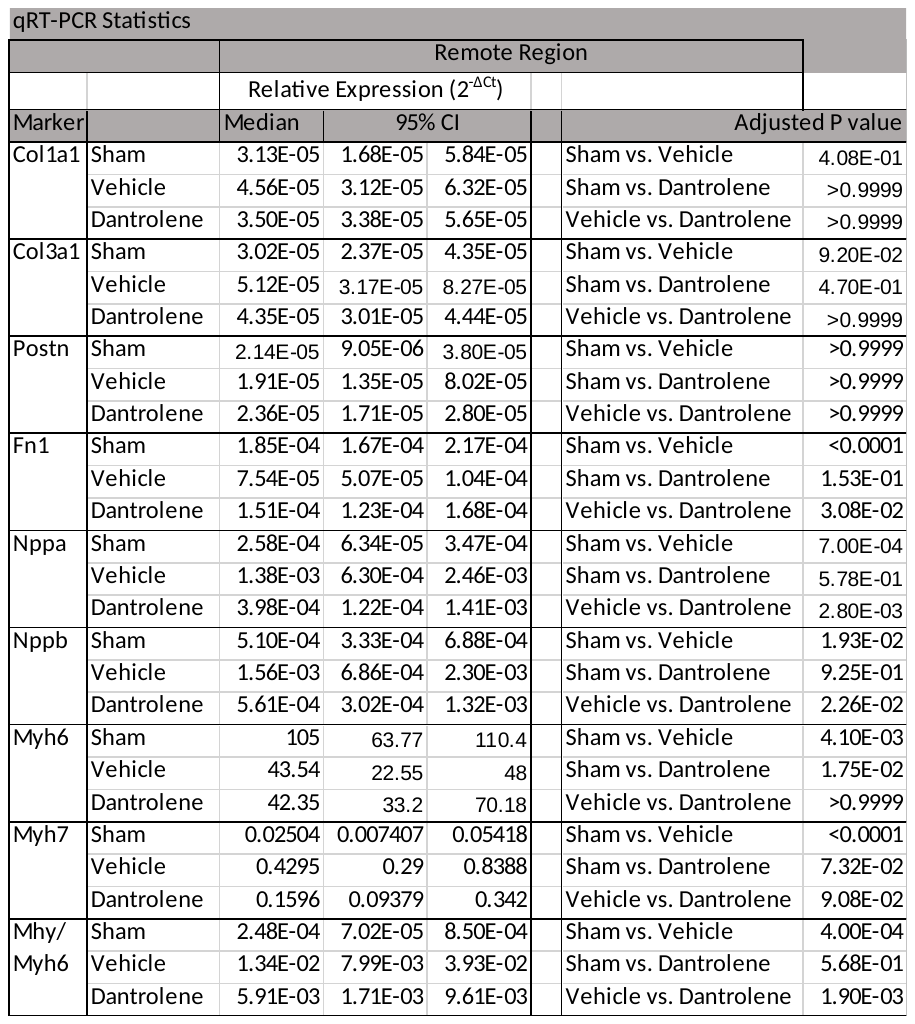

**Sup Table 2: qRT-PCR values from Border Zone and Remote Regions.** Relative gene expression to 18S rRNA content. [Sham = 6, Vehicle n=9, Dantrolene n=9]

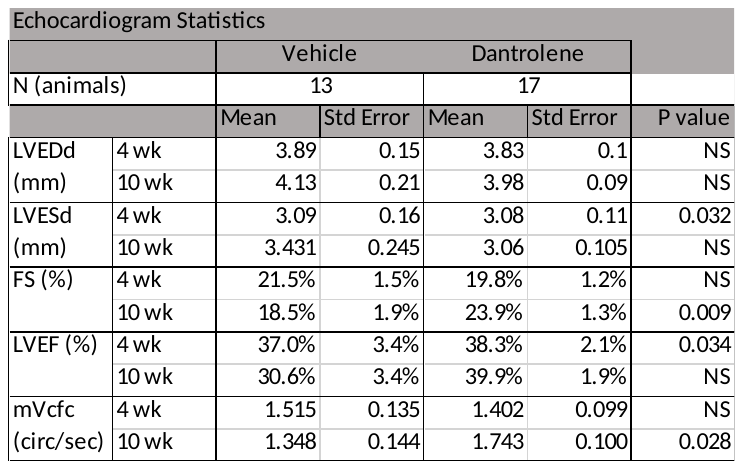

**Sup Table 3: Echocardiographic data.** [Sham = 6, Vehicle n=9, Dantrolene n=9]
